# Overexpression of UCP4 in astrocytic mitochondria prevents multilevel dysfunctions in a mouse model of Alzheimer’s disease

**DOI:** 10.1101/2022.01.25.477694

**Authors:** Nadia Rosenberg, Maria Reva, Leonardo Restivo, Marc Briquet, Yann Bernardinelli, Anne-Bérengère Rocher, Henry Markram, Jean-Yves Chatton

## Abstract

Alzheimer’s disease (AD) is becoming increasingly prevalent worldwide. It represents one of the greatest medical challenge as no pharmacologic treatments are available to prevent disease progression. Astrocytes play crucial functions within neuronal circuits by providing metabolic and functional support, regulating interstitial solute composition, and modulating synaptic transmission. In addition to these physiological functions, growing evidence points to an essential role of astrocytes in neurodegenerative diseases like AD. Early stage AD is associated with hypometabolism and oxidative stress. Contrary to neurons that are vulnerable to oxydative stress, astrocytes are particularly resistant to mitochondrial dysfunction and are therefore more resilient cells. In our study, we leveraged astrocytic mitochondrial uncoupling and examined neuronal function in the 3xTg AD mouse model. We overexpressed the mitochondrial uncoupling protein 4 (UCP4), which has been shown to improve neuronal survival *in vitro*. We found that this treatment efficiently prevented alterations of hippocampal metabolite levels observed in AD mice, along with hippocampal atrophy and reduction of basal dendrite arborization of subicular neurons. This approach also averted aberrant neuronal excitability observed in AD subicular neurons and preserved episodic-like memory in AD mice assessed in a spatial recognition task. These findings show that targeting astrocytes and their mitochondria is an effective strategy to prevent the decline of neurons facing AD-related stress at the early stages of the disease.

## Introduction

Alzheimer’s disease (AD) is the most prevalent cause of dementia in elderly people and is characterized by progressive deterioration of memory, language, and cognition. The number of cases is progressing worldwide in an epidemic fashion as humans tend to live increasingly longer (Alzheimer’s disease facts and figures, 2021). However, at this time, none of the medications available can slow down or stop the disease progression.

AD has been characterized by dysfunctions of energy metabolism, mainly glycolytic and mitochondrial functions, at early stages (Mosconi et al., 2008). A key function of mitochondria is ATP production via oxidative phosphorylation, which comes with the production of reactive oxygen species (ROS) as byproducts. The imbalance between the rate of ROS production and its clearance leads to oxidative stress, consistently reported in AD (Wang et al., 2014). The resulting changes in the oxidative environment are associated with glucose metabolism alteration, often by decreasing the activity of glycolytic enzymes (Butterfield & Halliwell, 2019; Mosconi et al., 2008). In addition, energy deprivation resulting from decreased oxygen and glucose delivery across blood vessels (Mosconi et al., 2008) renders neurons particularly vulnerable.

Mitochondrial dysfunction in astrocytes has been proposed to contribute to oxidative stress in AD, based on transcriptomic analyses of astrocytes in mouse models of AD and samples from patients (Abramov et al., 2004; Galea et al., 2022; Orre et al., 2014). Furthermore, ultra-deep proteomic analysis of cerebrospinal fluid (CSF) from patients with AD indicate that a large proportion of the dysregulated proteins are mitochondrial (Wang et al., 2020), many of them being highly abundant or mainly present in astrocytes (*see* http://www.brainrnaseq.org/). Because astrocyte mitochondria play a central role in the metabolic coupling between astrocytes and neurons (Vicente-Gutierrez et al., 2019), they may be important contributors to the bioenergetic breakdown observed in AD.

In previous studies, we found that increased expression of uncoupling protein 4 (UCP4) in primary mouse astrocytes leads to thorough and complex metabolic reorganization, reduced ROS production, and an increase in neuronal survival (Perreten Lambert et al., 2014a), possibly linked to the increased production of lactate. UCPs are proton carriers of the inner mitochondrial membrane that uncouple respiration from ATP production by partially dissipating the proton gradient generated by the electron transport chain, with the consequence of among other things diminishing mitochondrial ROS production (Ramsden et al., 2012).

In the present study, we focused on hypometabolism, appearing at preclinical stages before the onset of disease (Costantini et al., 2008; Mosconi et al., 2008). As neurons face oxidative stress already at early AD stages and as astrocytes present metabolic versatility compared to neurons, targeting astrocytic mitochondria could represent a valuable entry point to support the function of neurons facing AD-related stress. We investigated whether targeted overexpression of UCP4 in astrocytes mitochondria is neuroprotective in a mouse model of Alzheimer’s disease. We selected the triple transgenic AD (3xTg-AD) mouse model, which presents calcium dysregulations (Chakroborty et al., 2009), plasticity (Chakroborty et al., 2012) and cognitive impairment at young age (Clinton et al., 2007), before the onset of neuropathophysiological features (Oddo et al., 2003). In addition to preventing hippocampal atrophy, we show that astrocytic overexpression of UCP4 in the hippocampus of 3xTg mice preserved the dendritic architecture, prevented the development of aberrant electrophysiological properties of subicular neurons, and maintained the metabolite profile to wild-type levels. Most significantly, the treatment successfully preserved episodic-like memory in AD mice.

## Materials and Methods

### Animals

All the experimental procedures complied with the Swiss National Institutional Guidelines on animal experimentation and were approved by the canton of Vaud Cantonal Veterinary Office Committee for Animal Experimentation (authorization #VD3106 and VD3106_1). In all cases special attention was given to the implementation of 3Rs, housing, environmental conditions, and analgesia to improve the animals’ welfare. In all experiments, homozygous 3xTg mice (The Jackson Laboratory, MMRRC stock #34830, RRID:MMRRC_034830-JAX) (Oddo et al., 2003) adult male mice 7-10 months old were used and compared to age-and sex-matched WT (B6129SF2/J) mice, which are derived from the original mouse line with the same genetic background. We choose to work with males to avoid possible additional factors impacting AD, such as hormones affecting females (Clinton et al., 2007). In addition, males present at an early stage equal metabolic alterations as females (Dong & Brewer, 2019), rendering them suitable for our investigations.

### Viral vector construction

AAV plasmid pAAV-GfaABC(1)D-eGFP containing the minimal GFAP promoter (GfaABC(1)D) and the enhanced Green Fluorescent Protein (eGFP) between AAV inverted terminal repeats (ITRs) sequences were kindly provided by Dr. Bernard Schneider (Bertarelli Foundation Gene Therapy Core Facility, EPFL, Switzerland) and Dr. Sylvain Lengacher (GliaFarm SA, Campus Biotech, Switzerland) with the previously described DNA sequence (Dirren et al., 2014). A *de novo* synthesized mCherry-P2A-UCP4-HAHAHA-WPRE (GeneScript, New Jersey, USA) with the previously described mCherry (Shaner et al., 2004) and UCP4 (Perreten Lambert et al., 2014b) linked in frame with a 2A self-cleaving peptide sequence (P2A) was cloned into pAAV-GfaABC(1)D-eGFP, removing eGFP genes to obtain pAAV-GfaABC(1)D-mCherry-P2A-UCP4-HAHAHA-WPRE. pAAV-GfaABC(1)D-mCherry was obtained by cloning the mCherry gene from pAAV-GFAP-mCherry (Bernardinelli et al., 2014) in pAAV-GfaABC(1)D-mCherry-P2A-UCP4-HAHAHA-WPRE, removing mCherry-P2A-UCP4-HAHAHA genes. pAAV-GfaABC(1)D-UCP4-HAHAHA-WPRE was obtained by cloning UCP4-HAHAHA from pIRES-UCP4-HAHAHA (Perreten Lambert et al., 2014b) into pAAV-GfaABC(1)D-mCherry-P2A-UCP4-HAHAHA-WPRE, removing mCherry-P2A-UCP4-HAHAHA. AAV virus particles were generated as described by Grieger et al. (Grieger et al., 2006) at a titer ranging from 110^12^ to 110^13^ (Vg/ml), based on a recombinant genome containing the AAV2 ITRs and pseudotyped with an AAV capsid derived from serotype 9.

### Viral vector delivery and surgery

An initial dose of analgesic (Carprofen i.p. 5mg/kg) was administered subcutaneously to the 2-months-old mice before surgeries. Mice were anesthetized with isoflurane and placed on a heating blanket to maintain the body temperature at 37°C. The animal was head-fixed on a stereotaxic head frame apparatus (Stoelting, Wood Dale, IL). A small incision was made on the skin and the bone was exposed at the desired injection site. Viruses were injected with a thin glass pipette pulled on a vertical puller (Narishige, Tokyo, Japan). WT and 3xTg-AD mice were injected bilaterally in two sites in the hippocampus (site 1. Antero-posterior: -1mm, mediolateral: ±1.46mm, ventral: -1.5 from bregma; site 2. Antero-posterior: -1.56mm, mediolateral: ±2.15mm, ventral: -1.5 from bregma) with a virus encoding AAV9-mCherry or AAV9-mCherry-UCP4 (viral titer: 6.2 10^12^Vg/ml). 3µl was injected at a rate of 100-200nl/min in each injection site.

### In vivo MRI

MRI experiments were performed on a Bruker 3 Tesla small-bore scanner (BioSpec 30/18). A radiofrequency (RF) transmit-receive volume coil (Bruker BioSpin MRI, Ettlingen, Germany) with 82mm inner diameter and with active detuning was used for excitation, while a receive-only 2×2 channel mouse brain surface array coil (Bruker BioSpin MRI, Ettlingen, Germany) was used for signal detection. Animals were imaged six months after viral injections. They were isoflurane-anesthetized during the whole imaging session. The respiration rate was monitored through a respiration pad (SA Instruments, Stony Brook, NY, USA) placed against the mouse’s abdomen. The body temperature was monitored with a probe placed against the mouse flank and maintained at 37±0.5°C using a tubing system with circulating hot water. A fast low angle shot (FLASH) MRI pulse sequence with the following parameters: repetition time (TR)=100ms, echo time (TE)=7ms, RF excitation angle α=40°, field of view (FOV)=30×30mm, matrix size (MS)=256×256, slice thickness (ST)=1mm, number of slices=3, number of averages (NA)=2, acquisition time=32s, was used to acquire 3-plane localizer images. Then, turbo spin echo (TSE) T1-weighted images with the following parameters: TR=3300ms, TE=60ms, echo spacing=12ms, echo train length (ETL)=12, FOV=20×20mm, MS=128×128, ST=0.5mm, number of slices=19, NA=24, acquisition time=13min 12s, were acquired to visualize the hippocampus. MR image analysis was performed with Matlab. Hippocampi volumes were measured from multiple ROIs manually drawn by a single-blinded operator on the sagittal plane.

### Behavioral assay

Novel Object Location testing was performed using an arena consisting of a custom-built gray box (W=L=H: 45cm). The light source was placed below the horizon of the arena and provided 45 Lux, measured from the center of the arena. Extra lighting was provided by two short-range infrared illuminators (Raytec, Raymax RM25) mounted on the room ceiling. The objects were geometric shapes (W=L: 6.5cm and H: 4cm) built with Lego Duplo™. The color of the objects was uniform blue, red, or green and it was confounded across animals and groups. Mice were brought to the experimental room 30min before the beginning of the behavioral assay. The experiment consisted of 6 trials (duration: 5min, inter-trial interval: 3min) run in one single day (between 10:00 and 16:00).

In trial one, the habituation trial, each mouse was individually placed in the empty arena. During trials 2 through 4, the exploration trials, three objects were placed according to the pattern depicted in Fig. 5A (middle). These trials were used as an internal control to let the animals familiarize themselves with the objects and to identify potential bias towards one object over the others. Indeed, mice that showed more than 50% of contact toward one particular object in at least two sessions were excluded from further analyses. In trials 5 and 6, the novel object location (NOL) trials, one object was moved (object A) by 20cm to define the pattern depicted in Fig. 5A (right). An overhead camera (Basler, acA1300-60gm) driven by the Ethovision software (Noldus Information Technology) automatically tracked the mouse position across trials. The traveled distance was measured throughout the trials. An exploratory event directed toward the object was defined as the entrance of the mouse nose within a zone of 2cm from the object border; the object exploration zone (OEZ). The total number of entrances in the OEZ was used to calculate a preference index throughout sessions 2 to 6. Mice that did not show any contact with the objects during sessions 2 through 4 or that had a strong preference for one of the 3 objects (defined as >50% contacts directed toward one of the objects during 2 out of 3 trials of the exploratory trial) were excluded from subsequent analyses. Following these criteria definitions, a total of 26 subjects were excluded from the analyses. In **Fig. S6A** one outlier in the 3xTg-AD UCP4 group was detected but was not excluded from further analyses. Reaction to spatial change in the NOL trials was defined as the total number of contacts with object A across trials 5 and 6 and was expressed as % contacts over the total number of contacts with the 3 objects. The graphs report the % contacts with object A compared to the average % contacts with the two non-displaced objects (B and C). Behavior analyses were performed by an experimenter blinded to the treatment allocation of each experimental group.

### Immunohistochemistry and confocal microscopy

For virus testing, 4 weeks after injections, adult mice from both genotypes were transcardially perfused with cold 4% PFA (pH 7.4). Sagittal sections (30μm) were obtained using a vibratome (VT 1000 S, Leica). After patch-clamp recordings, acute brain slices (300µm thick) were post-fixed in 4% paraformaldehyde (PFA) for >24h. Then, the same procedure after both experiments was performed for immunostaining: slices were first washed with PBS and permeabilized with 1% Triton + 1g/l azide. Non-specific binding was blocked with PBS containing 2% normal horse serum. Samples were then incubated with the primary antibodies in a blocking solution for 5 days at 4°C: mouse anti-GFAP (G3893, Sigma, 1:300), mouse anti-HSP70 (MA3-028, Invitrogen, 1:500) and rabbit anti-HA (ab9110, Abcam, 1:200). Floating sections were incubated with respective secondary antibodies Alexa Fluor-488 goat anti-mouse (Invitrogen, 1:200), IRDye 680RD goat anti-rabbit (LI-COR, 1:200) and streptavidin Alexa Fluor 647 (Invitrogen, 1:500) overnight at 4°C. Finally, slices were incubated with DAPI (ThermoFisher, 1:1000) and mounted for imaging performed using Zeiss LSM780 and Zeiss LSM900 confocal microscopes.

### Metabolomic Profiling

#### Sample collection for metabolomic profiling

Focused Beam Microwave Irradiation was selected as euthanasia method as it optimally preserves metabolite levels after death by instantly inactivating enzymes of cellular metabolism (Matsui et al., 2017). Mice were first anesthetized with a mixture of Medetomidine (0.5mg/kg), Fentanyl (0.05mg/kg) et Midazolam (5mg/kg) through i.p. injection. They were then swiftly placed into the restrainer of a microwave fixation system (MMW-05 Muromachi, Tokyo, Japan). A pulse of 4kW with 0.6sec exposure was applied to cause the the immediate death. Mice were then rapidly decapitated, hippocampi dissected out, individually placed in plastic tubes, and stored on dry ice before tissue processing.

#### Metabolite extraction

Mouse hippocampus samples were pooled according to genotype and treatment. Samples were extracted by the addition of MeOH:H2O (4:1) (800μL). This solution containing hippocampal samples was further homogenized in the Cryolys Precellys 24 sample Homogenizer (2×20sec at 10000rpm, Bertin Technologies, Rockville, MD, US) with ceramic beads. The bead beater was air-cooled down at a flow rate of 110 L/min at 6 bar. Homogenized extracts were centrifuged for 15min at 4000g at 4°C (Hermle, Gosheim, Germany) and the resulting supernatants (100μL) were collected and evaporated to dryness in a vacuum concentrator (LabConco, Missouri, US). Dried sample extracts were re-suspended in MeOH:H_2_O (4:1, v/v) before LC-MS/MS analysis according to the total protein content (quantified using BCA protein Assay Kit (Thermo Scientific, Massachusetts, USA).

#### Multiple pathways targeted LC-MS/MS analysis

Tissue extracts were analyzed by Hydrophilic Interaction Liquid Chromatography coupled to tandem mass spectrometry (HILIC - MS/MS) in both positive and negative ionization modes using a 6495 triple quadrupole system (QqQ) interfaced with 1290 UHPLC system (Agilent Technologies). In positive mode, the chromatographic separation was carried out in an Acquity BEH Amide, 1.7μm, 100mm×2.1mm I.D. column (Waters, Massachusetts, US). The mobile phase was composed of A=20mM ammonium formate and 0.1% FA in water and B=0.1% formic acid in ACN. The linear gradient elution from 95% B (0-1.5min) down to 45% B was applied (1.5min-7min) and these conditions were held for 2min. Then initial chromatographic condition was maintained as a post-run during 5min for column re-equilibration. The flow rate was 400μl/min, column temperature 25°C and sample injection volume 2μl. In *negative mode*, a SeQuant ZIC-pHILIC (100mm, 2.1mm I.D. and 5μm particle size, Merck, Damstadt, Germany) column was used. The mobile phase was composed of A=20mM ammonium Acetate and 20mM NH_4_OH in water at pH 9.7 and B=100% ACN. The linear gradient elution from 90% (0-1.5min) to 50% B (8-11min) down to 45% B (12-15min). Finally, the initial chromatographic conditions were established as a post-run during 9min for column re-equilibration. The flow rate was 300μl/min, column temperature 30°C and sample injection volume 2μl. ESI source conditions were set as follows: dry gas temperature 290°C and flow 14 L/min, sheath gas temperature 350°C, nebulizer 45 psi, and flow 12l/min, nozzle voltage 0V, and capillary voltage +/-2000V. Dynamic Multiple Reaction Monitoring (dMRM) was used as the acquisition mode with a total cycle time of 600ms. Optimized collision energies for each metabolite were applied.

Pooled QC sample solution (representative of the entire sample set) was analyzed periodically (every 5 samples) throughout the overall analytical run. In addition, a series of diluted quality controls (dQC) were prepared by dilution with methanol: 100% QC, 50% QC, 25% QC, 12.5% QC and 6.25% QC and analyzed at the beginning and at the end of the sample batch.

#### Data (pre)processing and quality assessment

Raw LC-MS/MS data were processed using the Agilent Quantitative analysis software (version B.07.00, MassHunter Agilent technologies). Relative quantification of metabolites was based on EIC (Extracted Ion Chromatogram) areas for the monitored MRM transitions. The exported data (comprising peak areas of detected metabolites across all samples) were corrected for the signal intensity drift (over time) using LOWESS/Spline algorithm. The metabolite features represented by “not-well behaving” peaks (CV (QC peaks) > 30% & R^2^ (QC dilution curve) were discarded from further statistical analysis.

#### Statistical data analysis

Two-way Anova (on log10 transformed peak abundances) was used to test the significance of metabolite changes between different groups of samples (3xTg-UPC4, 3xTg, WT-UPC4 and WT) with an arbitrary level of significance, p-value = 0.05 (and adjusted p-value corrected for multiple testing with Benjamini-Hochberg method).

#### Acute slice preparation, patch-clamp recordings

Mice were deeply anaesthetized with ketamine (100mg/kg, ip) and xylazine (10mg/kg, ip) and then transcardially perfused with 10-20ml of ice-cold NMDG-based artificial cerebrospinal fluid (ACSF) containing (mM): 93 NMDG, 2.5 KCl, 1.2 NaH_2_PO_4_, 30 NaHCO_3_, 20 HEPES, 25 glucose, 5 ascorbic acid, 2 thiourea, 3 sodium pyruvate, 12 N-acetyl-L-cysteine, 10 MgSO_4_, 0.5 CaCl_2_ (Martinez-Losa et al., 2018), pH 7.3, equilibrated with 95% O_2_ and 5% CO_2_. Next, mice were decapitated and their brains quickly removed and placed into the same NMDG ACSF. 300µm thick sagittal acute slices were prepared using a vibratome (VT1000S, Leica) in the same ice-cold ACSF. Slices were incubated in ACSF containing (mM): 92 NaCl, 2.5 KCl, 1.2 NaH_2_PO_4_, 30 NaHCO_3_, 20 HEPES, 25 glucose, 5 ascorbic acid, 2 thiourea, 3 sodium pyruvate, 12 N-acetyl-L-cysteine, 2 MgSO_4_, 2 CaCl_2_, for 20min at 34°C and then kept at room temperature for at least 20min before being transferred to the recording chamber. Slices were held down by a metal harp and superfused by oxygenated ACSF containing (in mM): 119 NaCl, 2.5 KCl, 1 NaH_2_PO_4_, 26.2 NaHCO_3_, 11 glucose, 1.3 MgSO_4_, 2.5 CaCl_2_ at 32°C. Recordings were made under a Zeiss LSM510 Meta upright microscope equipped with infrared DIC using a 40X water-dipping objective lens (Carl Zeiss). Whole-cell patch-clamp of subicular neurons was obtained with borosilicate glass pipettes (6-8 MΩ resistance). The patch-clamp intracellular solution contained (in mM): 130 K-gluconate, 5 NaCl, 5 KCl, 1 MgCl_2_, 0.1 EGTA, 0.025 CaCl_2_, 10 HEPES, 4 glucose, 5 Na phosphocreatine, 4 Mg-ATP, 0.3 Na-GTP, pH 7.3 with KOH, 290 mOsm. 0.1% biocytin was added to the pipette solution to further visualize the neurons patched. Recordings were obtained in current-clamp configuration using a Multiclamp 700B amplifier (Molecular Devices). Data were acquired at 10kHz and filtered at 2kHz (Digidata 1440 analog-to-digital converter), controlled with the pCLAMP 10 software. Experiments were discarded if the access resistance, monitored by -5 mV steps (0.1Hz), varied by more than 20% or a holding current at -70mV larger than -150 pA. Current steps (30pA increments) of 320ms duration were injected to measure burst frequency and of 2000ms for assessing AHPs. In a subset of experiments, the SK channel enhancer ns-309 (5µM, Sigma-Aldrich) was bath applied 10min before repeating the same measurements. The amplitude of AHPs and burst frequency were quantified using the Clampfit software.

#### Sholl and morphometric analysis

Morphological analyses of neurons were made from 80-100μm confocal image stacks acquired using a 20X 0.8NA Plan Apochromat objective lens (Carl Zeiss) on a Zeiss LSM900 microscope, and 3D-reconstructed using Imaris Filament Tracer (Bitplane AG, Switzerland). Reconstructions were performed in a blinded experimental design. For each reconstructed 3D morphology, we performed a Sholl and morphometry analysis (Sholl, 1953) for the following dendritic compartments: basal, apical and whole dendritic tree. For Sholl analysis, concentric rings were placed on the cell-centered at the soma. The radius of the ring was incremented by 5µm steps until dendrites were completely covered. Number of dendritic crossings with the ring was calculated for each ring radius. For morphometry analysis, we measured the number and length of dendritic branches, using open-source Python package NeuroM (https://github.com/BlueBrain/NeuroM).

#### Electrophysiological modeling

The features from voltage recordings were extracted using the BluePyEfe package (https://github.com/BlueBrain/BluePyEfe) and Electrophys Feature Extraction Library (eFEL, https://github.com/BlueBrain/BluePyEfe). For each cell a rheobase was calculated. Features were extracted from the recordings of a chosen fraction from rheobase and averaged among the sweeps and cells. Number of spikes, inter spikes intervals, voltage base, AHP after each spike, time of AHP relative to AP peak, first second and last AP amplitudes, the voltage at the beginning of AP, slope and coefficient of variation (CV) of interspike intervals, AP width and amplitude of the AHP after the stimuli were extracted from voltage recordings in response to depolarizing steps (for 200ms step we considered step amplitudes of 200, 300 and 500% from the rheobase, for 2000ms step -100, 150, 300 and 400% from the rheobase). Base voltage, voltage deflections, and voltage at the steady-state after the stimuli were extracted for the hyperpolarization step.

The neuronal model consisted of the 3D morphological reconstruction and several ionic mechanisms. The following mechanisms were considered: calcium high voltage-activated channels (Reuveni et al., 1993), calcium low voltage-activated channels (Avery & Johnston, 1996), persistent and transient potassium currents (Korngreen & Sakmann, 2000), SK type calcium-activated potassium current (Köhler et al., 1996), Kv1.3 potassium current (Rettig et al., 1992), Ih current (Kole et al., 2006), transient and persistent sodium currents (Colbert & Pan, 2002) and intracellular calcium dynamic (Destexhe et al., 1994). The models of the mechanism are adapted Markram et al. (Markram et al., 2015). The maximal conductance of the aforementioned mechanisms, leak conductance, membrane capacitance, leak reversal potential and exponential decays of sodium and Ih current were set as free parameters, while axial resistance was set to 100 Ohm/cm. Initial simulation voltage was set to − 80 mV, simulation temperature to 34°C, and equilibrium potentials for sodium and potassium to 50 mV and − 90 mV, respectively. All parameters of ionic mechanisms, such as membrane capacitance and intracellular calcium dynamics were specifically optimized for soma and dendrite, and axon. EPSP stimuli (Markram et al., 2015) were placed in the middle of the morphological compartment.

The models were implemented in NEURON (Hines & Carnevale, 1997), and optimization of ionic maximal conductance and calcium dynamics parameters were performed with IndicatorBased Evolutionary Algorithm (IBEA) using the BluePyOpt python package (Van Geit et al., 2016). A successful model was considered to have all feature errors less than 5 SD of the experimental mean feature. We generated 100 models for each group of neurons, using different seed values, to diversify the output. Several models with the best standard scores were chosen (WT n=5, WT UCP4 n=5, 3xTg n=7, 3xTg-AD UPC4 n=5).

#### Statistics and data analysis

The statistical tests used are listed in the respective Figure legends. Graphs and statistical analyses were performed with GraphPad Prism.

## Results

### Selective UCP4 expression in mitochondria of hippocampal astrocytes

The atrophy of the subicular region is the earliest hippocampal anatomical marker of AD (Carlesimo et al., 2015). The subiculum receives inputs from CA1, which itself receives inputs from CA3 (van Strien et al., 2009). We therefore hypothesized that preserving the neuronal function of the hippocampal network and subiculum will protect mice against AD-related memory impairment. Because UCP4 overexpression in astrocytes has neuroprotective effects *in vitro* (Perreten Lambert et al., 2014b), we postulated that *in vivo* overexpression of UCP4 would have beneficial effects on neurons facing AD. To elevate the levels of UCP4 in astrocytes, we designed adeno-associated viruses (AAV) that specifically target mitochondria of astrocytes, namely a vector expressing mCherry only under the GfaABC(1)D astrocytic promoter, which was used as a control, and a vector expressing mCherry coupled to UCP4 (Fig. 1A). A poly HA-tag was added to the UCP4 sequence for later accurate detection of vector expression by immunohistochemistry. The inserted 2A self-cleaving peptide sequence allows yielding the same level of expression of mCherry and UCP4. After testing different AAV capsids, we selected serotype 9 which was found to be most efficient at specifically inducing astrocytic expression in the hippocampus. Viral vectors were injected in the dorsal CA1-CA3 and subicular regions of the hippocampus (Fig. 1C). We injected the viral vectors into four separate groups: two groups of wild-type (WT) and two groups of 3xTg-AD mice. These groups are referred to as WT mCherry (WT), WT mCherry-UCP4 (WT UCP4), 3xTg-AD mCherry (3xTg), and 3xTg-AD mCherry-UCP4 (3xTg UCP4). Thus, mice of all groups were injected either with the control mCherry or the mCherry-UCP4 encoding virus at two months of age (Fig. 1B), before overt neuropathology, mitochondrial impairment, memory impairment, or synaptic dysfunction (Oddo et al., 2003).

**Fig. 1.**
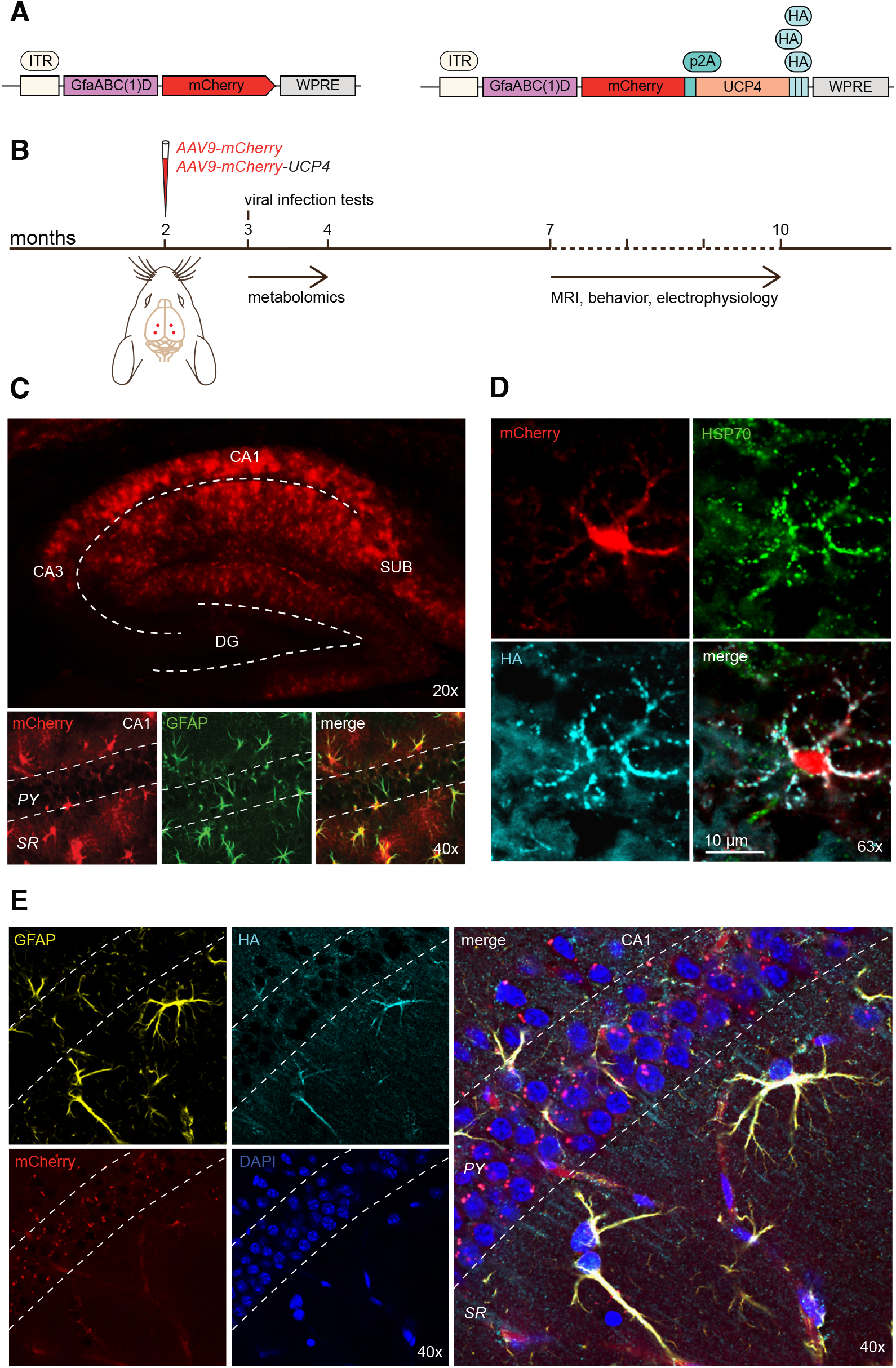
Characterization and validation of viral vectors used to drive UCP4 expression in astrocytes of the hippocampus. (**A**) Design of AAV vectors used to infect astrocytes. *Left*: construct containing mCherry alone under astrocytic promoter (GfaABC(1)D) used as a control. *Right*: Vector containing mCherry coupled to UCP4, which contains a poly-HA tail. (**B**) Timeline of experimental procedures. Two-months old WT and 3xTg-AD mice were injected bilaterally in the hippocampus with the AAV9-mCherry or the AAV9-mCherry-UCP4 viral vectors (red dots: injection sites). Animals were tested starting at 7 months of age. (**C**) Confocal images showing mCherry expression (red), GFAP staining (green) and the merged image in the hippocampus one month after viral infection (DG; dentate gyrus, SUB: subiculum). Images show that mCherry colocalizes with GFAP positive astrocytes (green). (**D**) Immunostaining of the mitochondrial marker HSP70 (green) and UCP4 HA-tag (cyan) plus mCherry expression (red) and the merged image, showing that tagged UCP4 is expressed in mitochondria of astrocytes. (**E**) Immunostaining of GFAP (yellow) and HA-tag (cyan) plus mCherry indicates that tagged UCP4 is still expressed 10 months post-injection and remains present in astrocytes. DAPI staining (blue) was used to visualize the CA1 pyramidal layer. (*PY*: pyramidal layer, *SR*: stratum radiatum).

Fig. 1C confirms that one-month post-injection, mCherry-infected cells are astrocytes by colocalizing with glial fibrillary acidic protein (GFAP) immunostaining. We evaluated the mitochondrial expression of UCP4 using HA-tag immunostaining, which colocalized with the mitochondrial protein HSP70 in astrocytes (Fig. 1D). As neurons were found not to be infected (*SI Appendix*, Fig. S1), we could conclude that the viral vectors enable selective expression of UCP4 in mitochondria of astrocytes. Furthermore, we found that UCP4 was still expressed and colocalized with astrocytic mitochondria in 10 months old mice (Fig. 1E), when magnetic resonance imaging (MRI), behavioral, and electrophysiological experiments were carried out.

### Astrocytic UCP4 overexpression prevents metabolic alterations in 3xTg-AD hippocampi

After ensuring that UCP4 was still expressed in astrocyte mitochondria 10 months post-injection, we investigated how the metabolism of 3xTg mice was altered at the early stages of the disease and what was the effect of UCP4 overexpression. Multiple pathways targeted metabolite screen of central carbon metabolism (including 520 metabolites combining positive and negative ionization mode) revealed that a set of metabolites depicted in Fig. 2A was significantly upregulated in 3xTg mice compared to both WT groups at 3-4 months of age, when mice were still asymptomatic. Interestingly, the metabolic profile of 3xTg UCP4 hippocampi appeared similar to that of WT groups, consistent with a normalizing effect of the UCP4 treatment (Fig S2). Moreover, pathway analysis highlighting affected biochemical pathways associated with these metabolic changes, unveiled alterations in the TCA cycle, glycolysis and in the metabolism of amino acids (Fig. 2B). These results indicate that (a) profound metabolic perturbations occur in 3xTg hippocampi before overt AD symptoms, which are prevented in 3xTg UCP4 mice, and (b) that UCP4 overexpression in mitochondria of astrocytes is rapidly effective, as its effects appear already 1-2 months post-injection.

**Fig. 2.**
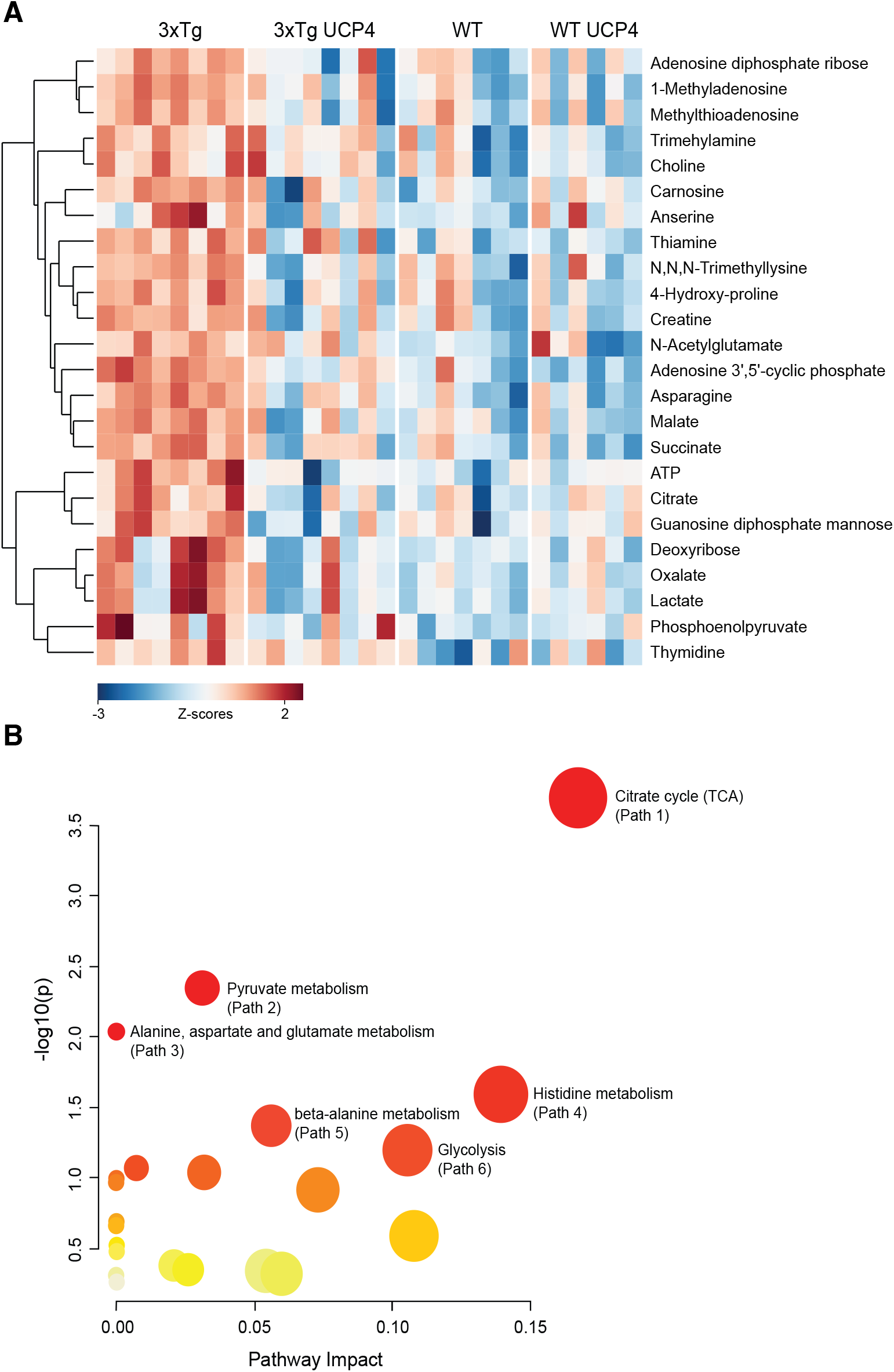
Astrocytic UCP4 overexpression maintains normal hippocampal metabolite profile in 3xTg-AD mice. (A) Heat map of metabolites whose levels changed significantly in 3xTg mice compared to both WT groups and 3xTg UCP4 mice (see also Fig. S2). Each row represents a metabolite and each column a different mouse. (B) Pathway analysis performed on metabolomics results showed in (A). Each node represents an altered metabolic pathway in the hippocampus of 3xTg mice that is preserved in 3xTg UCP4 mice and their size indicates the impact of this pathway.

### Hippocampal atrophy and dendritic shrinkage of subicular neurons occurring in 3xTg-AD mice are prevented by astrocytic UCP4 expression

Hippocampal atrophy is one of the most validated and extensively used biomarkers in AD (Apostolova et al., 2006). To investigate whether it is influenced by astrocytic UCP4 overexpression, we performed MRI and compared hippocampal volumes in each group of mice at 8 months of age. We found that 3xTg mice exhibited significant hippocampal atrophy (−18.6±5.5%) compared to WT groups (*P* = 0.001) (Fig. 3A). However, 3xTg UCP4 mice showed similar hippocampi volume as both WT groups and was significantly different from 3xTg-hippocampi (WT UCP4 vs 3xTg: *P* = 0.023; 3xTg UCP4 vs 3xTg: *P* = 0.046) (Fig. 3B). These results indicate that UCP4 overexpression prevents hippocampal atrophy.

**Fig. 3.**
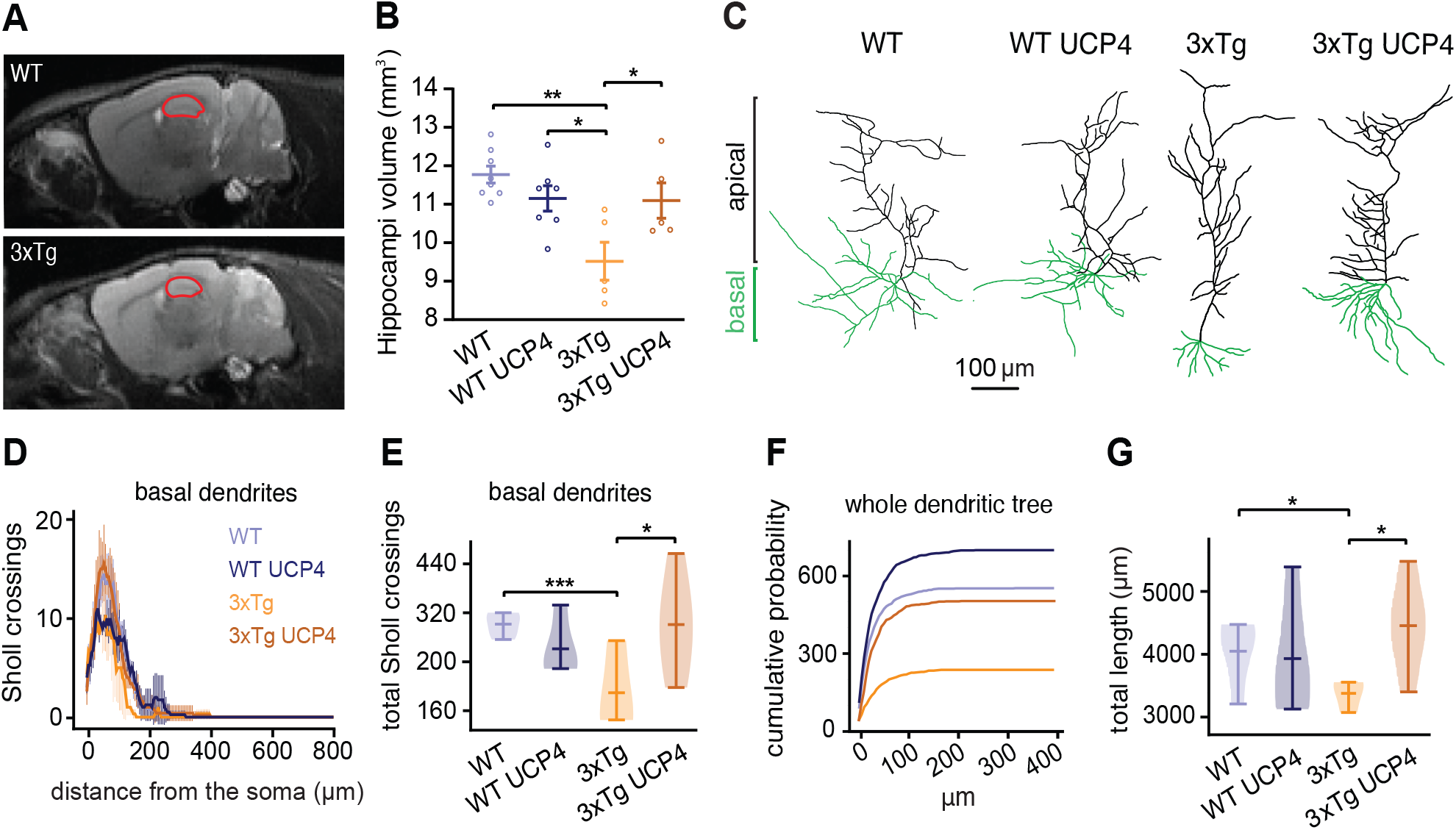
Hippocampal atrophy and alteration of dendritic arborization of 3xTg-AD neurons is impeded by UCP4 overexpression. (**A***) In vivo* MRI of mice hippocampus. Representative brain T2-weighted images of a WT and a 3xTg mouse in sagittal plane. Red dashed lines indicate ROI for the hippocampus. (**B**) Quantification of hippocampal volume (WT: n=8, WT UCP4: n=7, 3xTg: n=5, 3xTg UCP4: n=5). Each circle represents the volume of combined left and right hippocampus per mouse. (**C**) Examples of reconstructed morphologies from the 3D models with identified apical (black) and basal (green) dendrites. (**D**) Sholl profiles of basal dendrites, calculated using 5 μm increments (WT: n=4, WT UCP4: n=4, 3xTg: n=3, 3xTg UCP4: n=5). (**E**) Total number of Sholl crossings for each group. (**F**) Cumulative probability plot of total dendritic length (µm) for each group. (**G**) Total path length of dendritic tree for each group calculated per morphology. In (**B**) one-way ANOVA followed by Tukey’s post hoc test was performed (*F*_(3, 21)_ = 6.81). Error bars in this figure are mean ± SEM. \**P* < 0.05, \*\**P* < 0.01. In (**E** and **G**) nonparametric bootstrapping test was performed, \**P* < 0.05, \*\*\**P* < 0.001. Bars on violin plots: mean ± CI.

Loss of dendritic architecture integrity, which may partly contribute to cerebral atrophy, is observed in AD in a wide population of neurons, such as subicular pyramidal cells (Flood, 1991). As the subiculum represents the main output from the hippocampus and because it plays a pivotal role in coding spatial information (Kim et al., 2012), which is compromised in AD, we analyzed the dendritic tree of biocytin-filled subicular neurons. Z-stack fluorescence images of neurons were acquired for the reconstruction of their 3D structures (Fig. S3 A), which were subsequently used for Sholl and morphometric analysis (Fig. 3, D-G). We divided basal and apical processes to compare the dendritic fields among groups of neurons (Fig. 3C). 3xTg neurons displayed different dendritic arborization when compared to both WT groups or 3xTg UCP4 neurons. This observation was confirmed by Sholl’s analysis. Indeed, while apical dendrites did not present differences among groups (*SI Appendix*, Fig. S3 B and C), basal dendrites of 3xTg mice exhibited a significant decrease in Sholl crossings compared to WT mice (3xTg vs WT: *P*=0.001), corresponding to a marked reduction of the dendritic basal tree complexity (Fig. 3D). Interestingly, we found that UCP4 treatment of 3xTg mice maintained basal dendritic arborization to WT values (3xTg vs 3xTg UCP4: *P*=0.03) (Fig. 3E). A drastic reduction in dendritic length was revealed in 3xTg mice compared to both WT groups 3xTg vs WT: *P* = 0.02, 3xTg vs 3xTg UCP4: *P* = 0.04). However, this alteration was not observed in 3xTg mice overexpressing UCP4 (Fig. 3G). The same trend was observed when considering the total number of dendritic branches (Fig. S3 D). Thus, UCP4 overexpression in astrocytes conserves dendritic arborization of 3xTg subicular neurons.

### Astrocytic UCP4 overexpression prevents the development of aberrant electrophysiological properties in 3xTg-AD subicular neurons

The elaborate dendritic geometry of a neuron affects its physiological properties, such as its excitability (Rall, 1962; Šišková et al., 2014). For this reason, we decided to characterize the intrinsic excitability of subicular neurons by performing patch-clamp recordings in WT, 3xTg and the respective UCP4-treated groups (Fig. 4A). Resting membrane potential (RMP) (*P* = 0.248), membrane threshold to evoke a burst (*P* = 0.065), half -width (*P* = 0.115), rheobase (*P*=0.131), tau (*P*=0.973), input resistance (*P*=0.1) and sag (*P*=0.153) did not show differences between groups (Fig. S4, A-G). However, the membrane capacitance of WT and WT UCP4 neurons was found to be significantly higher than in 3xTg mice (WT vs 3xTg: *P* = 0.004; WT UCP4 vs 3xTg: *P* = 0.009), while the capacitance in 3xTg UCP4 group was not significantly different from the WT groups (Fig. S4 H). These results show that the viral vectors did not affect the general electrical properties of subicular neurons and that UCP4 overexpression maintains membrane capacitance of 3xTg mouse neurons close to values of WT subicular cells.

**Fig. 4.**
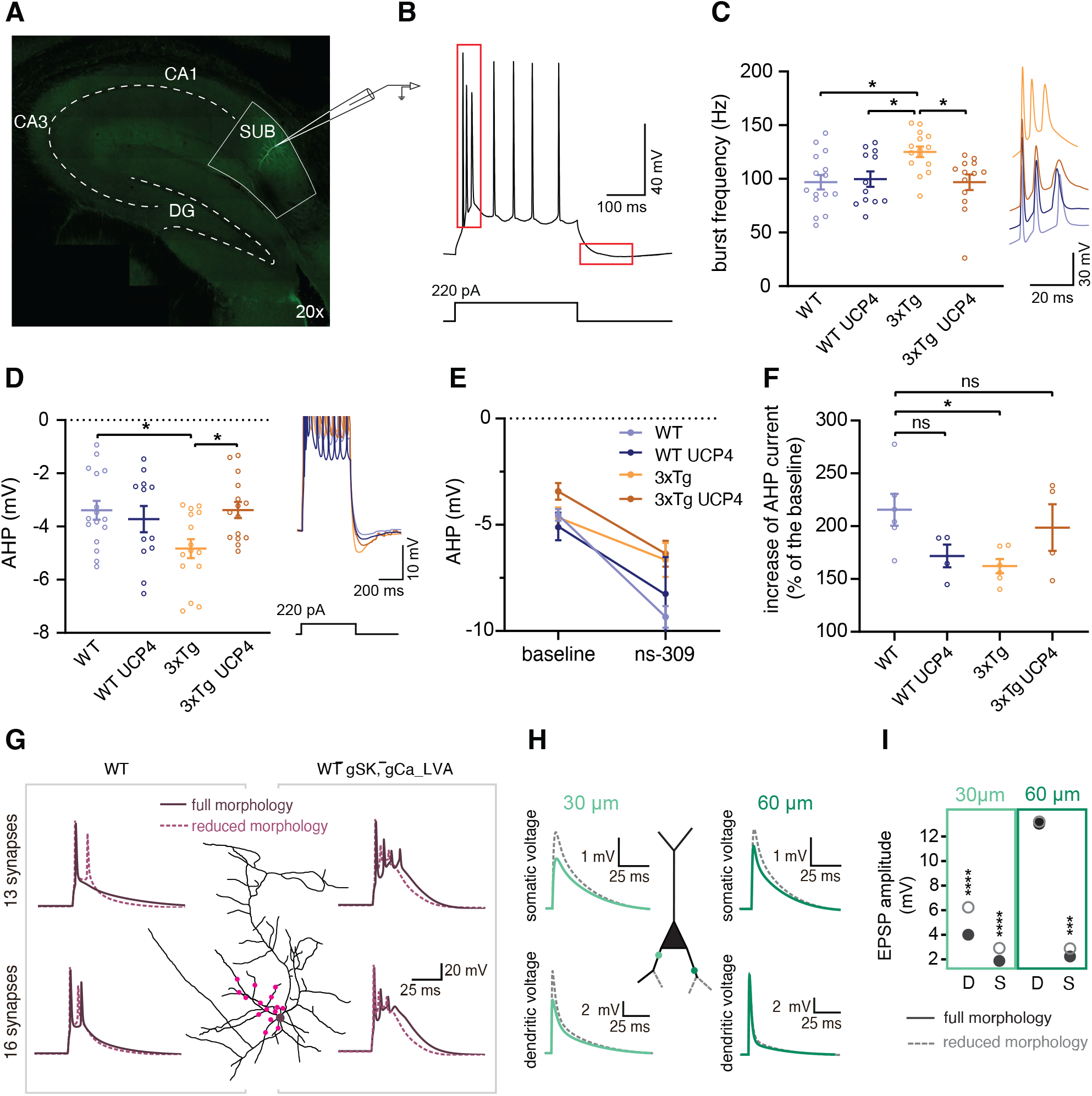
Hyperexcitability in 3xTg-AD subicular neurons is hampered by UCP4 overexpression and modeling makes the link between dendritic morphology and neuronal excitabiliy. (**A**) Confocal image of the hippocampus and the area of patch-clamp recordings highlighted by a white line (DG: Dentate gyrus, SUB: subiculum). (**B**) Example trace of a current-clamp recording. Red rectangles represent the temporal windows used to analyze the first burst frequency and the amplitude of AHP (**C** and **D**, respectively). (**C**) Plot of fire frequency of the first burst. Symbols represents the average of all the sweeps per cell. (WT: n=15, WT UCP4: n=12, 3xTg: n=15, 3xTg UCP4: n=13). *Inset*: representative traces of the first burst occurring after depolarization. (**D**) Amplitude of the spike-evoked AHP. Symbols represent the average AHP of all the sweeps per cell. (WT: n=16, WT UCP4: n=12, 3xTg: n=15, 3xTg UCP4: n=15). *Inset*: representative spike trains and subsequent AHP. (**E**) AHP amplitude in control condition and after SK channel activation using ns-309. Symbols represent the average of all mice. (WT: n=6, WT UCP4: n=4, 3xTg: n=6, 3xTg UCP4: n=4). (**F**) Plot showing the ns-309 mediated enhancement of AHP amplitude as a percentage of the baseline. Each circle represents one neuron (same n as in **E**). (**G**) *Center*: illustration of stimulation sites at different locations along the dendritic tree (pink dots) to elicit somatic spike in the soma (purple dot). *Left*: voltage traces when 13 EPSPs (top) or 16 EPSPs (bottom) are simultaneously injected in WT condition. *Right*: voltage traces when 13 EPSPs or 16 EPSPs are simultaneously injected in simulated increased somatic 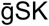 and dendritic 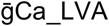. (**H**) Effect of morphology on signal propagation. Voltage traces in the soma (top traces) and in the dendrite (bottom traces) after EPSP stimulation at 30µm (light green rectangle) and 60µm (dark green rectangle) from the soma. In **G** and **H** Solid line: full WT dendritic length, dashed line: shortened dendritic tree that matches the 3xTg morphometrics analysis. (**I**) Quantification of panel **H** results: voltage amplitudes in dendrite (D) and soma (S) followed by EPSP stimulation for 4 WT models (same as in **Fig.3**). Dendritic and somatic amplitudes are statistically different in case of proximal EPSP stimulation (bootstrap permutation test, dendritic: *P*=0; somatic: *P*=0), while for distant stimulation statistical difference is only detected in somatic voltage. In (**C**) a Kruskal-Wallis followed by a Dunn’s post hoc test was performed, in (**D**) and (**F**) one-way ANOVA (**D**: *F*_(3, 54)_ = 3.536; **F**: *F*_(3, 16)_ = 3,475) followed by Tukey’s post hoc test was performed. Error bars in this figure are mean ± SEM. \**P* < 0.05, ns; not significant. In (**I**), bootstrap permutation test was performed. *** *P* < 0.001; **** *P* < 0.0001. Bars on violin plots: mean ± CI.

Next, we measured the instantaneous frequency of the first action potential (AP) burst (Fig. 4B) in response to each suprathreshold current step injection. Fig. 4C shows that 3xTg mice exhibit a significantly higher burst frequency elicited by current injection steps compared to WT (*P* = 0.017) and WT UCP4 mice (*P* = 0.049), pointing to increased excitability that was prevented in 3xTg UCP4 subicular neurons (3xTg vs 3xTg UCP4: *P* = 0.028). Depolarization-induced burst firing in subicular neurons is mediated by low-voltage-activated T-type calcium channels (Joksimovic et al., 2017), which participate in the increases in intracellular calcium. Cytosolic calcium plays a crucial role in neuronal excitability and limits firing frequency through SK channel activation. Because SK channels are small-conductance calcium-activated potassium channels underlying after-hyperpolarization potential (AHP) (Stocker et al., 1999), we examined AHP amplitude (Fig. 4B) after a train of bursts and APs generated by current step injections. The amplitude of the spike-evoked AHP was significantly larger in 3xTg neurons compared to WT (P = 0.0320). However, AHP amplitude in 3xTg UCP4 neurons presented the same pattern as both WT groups (3xTg vs 3xTg UCP4: *P* = 0.0344) (Fig. 4D). We hypothesized that if SK channels activity is enhanced in AD mice (Chakroborty et al., 2012) (Chakroborty et al., 2012), they should show occlusion upon activation, *i.e*. a strong AHP increase should be induced in WT neurons but not in 3xTg cells. To test this idea, we bath-applied ns-309 (5µM, 10min), a potent enhancer of SK channels (Nam et al., 2017), which increased AHP in the four groups (Fig. 4E). However, when normalized to the baseline, we found that SK-mediated AHP was significantly smaller in 3xTg mice compared to WT mice (*P* = 0.024). Interestingly, upon SK channel activation, 3xTg neurons treated with UCP4 showed an AHP increase comparable to that of WT neurons (WT vs 3xTg UCP4: *P* = 0.749; WT vs WT UCP4: *P* = 0.112) (Fig. 4F). These data indicate that the higher excitability and activity of SK channels found in 3xTg cells were kept to control levels by astrocytic UCP4 overexpression in 3xTg mice.

To investigate how morphological and electrical alterations shape spike emission and signal propagation after synaptic stimulation, we implemented the 3D morphological reconstructions (Fig. S3 A) and electrophysiological features extracted from patch-clamp recordings (Fig. 4) into full morphology models. Ionic mechanisms present in cells were defined and optimized such that the responses of the models reproduce features extracted from experiments (fig. S5 A). After model optimization for each of the four groups, simulations of 200, 300 and 450 pA current injections were produced to test how burst frequency and AHP would behave in the models. Current injections led to statistical differences between 3xTg and the other groups translated into an increase in burst frequency (3xTg vs WT: *P* = 8e-3; 3xTg vs WT UCP4: *P* = 5e-3; 3xTg vs 3xTg UCP4: *P* = 3e-3) and AHP (3xTg vs WT: *P* = 1e-3; 3xTg vs WT UCP4: *P* = 1e-3; 3xTg vs 3xTg UCP4: *P* = 1e-2) (fig. S5, B-C), as was observed in experimental data (Fig. 4, C-D). To simulate ex-vivo conditions of SK channels activation with bath applied ns-309 (5µM), SK channel conductance 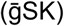 was manually increased and pooled across responses of the models to 200 and 450 pA current injections. We observed that in all groups the simulated 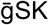 increase enhanced AHP compared to the baseline (fig. S5 D). The relative AHP increase was however significantly smaller in 3xTg models compared to other groups (3xTg vs WT: *P* = 1e-4; 3xTg vs WT UCP4: *P* = 0; 3xTg vs 3xTg UCP4: *P* = 5e-3) (fig. S5 E), as previously observed in electrophysiological recordings (Fig. 4F). Thus, the generated models reproduced reliably and accurately experimental data (Fig. S5, A-E).

### Computational modeling of neuronal properties supports the effect of SK channels and links truncated dendritic arborization with increased excitability of 3xTg neurons

SK channels a play crucial role notably in intrinsic excitability and in shaping postsynaptic responses (Ngo-Anh et al., 2005). Their activity is increased in 3xTg mice(Chakroborty et al., 2012). SK channels are voltage-independent channels, solely activated by elevations of intracellular calcium concentrations that may derive from different sources (Adelman et al., 2012). To identify key elements underlying the prevention of aberrant burst frequency and AHP by UCP4 overexpression, we selected a series of model parameters based on their relevance for cell excitability and SK channel activity (Fig. S5 F). Distribution analysis of these parameters unveiled a significantly higher conductance of SK channels in the 3xTg model compared to the other groups, a smaller conductance of low voltage-activated calcium (LVA) channels and an alteration of the percentage of free calcium in the soma. In dendrites, LVA presented the opposite behavior, having a higher conductance in 3xTg cells compared to simulated WTs and 3xTg UCP4 neurons (Fig. S5 F). We then used the models to test the impact of altered dendritic architecture on the output signal of subicular neurons. To do so, 13 or 16 excitatory postsynaptic potentials (EPSPs) were simultaneously injected at different locations of the dendritic tree to elicit somatic spiking (Fig. 4G, middle scheme) in WT model with intact dendritic tree (full morphology) and in models where dendrites were manually shortened by 24% to match the 3xTg morphometry (reduced morphology). Activation of 13 synapses elicits one spike, which turns into a burst in the reduced morphology (Fig. 4G, top left). If more synapses (16) are activated, the voltage response is converted into a burst, whose frequency is increased in the reduced morphology (Fig. 4G, bottom left). The same stimulations were then applied to full and reduced morphologies, with the additional implementation of increased somatic SK channels and dendritic low voltage-activated calcium (LVA) conductance 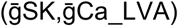 to fit the 3xTg model. Both activation of 13 and 16 synapses induce a burst that is of higher frequency in neurons with shorter dendrites (Fig. 4G, right). These results show that (a) a reduced dendritic arborization increase excitability in WT and WT 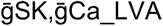 models, and (b) that in the WT 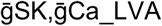 model, excitability is higher, and has a higher propensity of bursting.

The efficiency of signal propagation from dendrites to soma is a key factor for the induction of synaptic plasticity (Stuart & Spruston, 2015), which in turn is crucial for storing engrams in the brain (Takeuchi et al., 2014). We therefore investigated how voltage signals are propagated along the dendritic branches (Fig. 4H). When stimulated 30µm away from the soma, dendritic and somatic voltages show significantly higher EPSP amplitudes in reduced morphologies (dendritic: *P* = 0; somatic: *P* = 0), whereas at 60µm, only the somatic voltage trace displays a significantly higher amplitude in the cut morphology (Dendritic: *P* = 0.335; Somatic: *P* = 3e-3) (Fig. 4I). These results show that a reduced architecture of dendrites as observed in 3xTg mice, alters signal propagation towards the soma providing a causal link between impaired dendritic morphology and increased excitability.

### Prevention of spatial memory impairments by UCP4 overexpression in astrocytes

Hippocampal-dependent impairment of learning and memory has been linked to aberrant calcium signaling (Chakroborty et al., 2009) and cellular excitability in AD mice (Davis et al., 2014). The elevated firing rate of subiculum neurons is involved in spatial navigation (Kitanishi et al., 2021), a cognitive function jeopardized in AD patients. As memory for object location is largely hippocampus-dependent and is impaired in Alzheimer’s patients at the early stages of the disease (Kessels et al., 2010) we performed a novel object location task as depicted in Fig. 5A. During the habituation trial (Fig. 5A, *left*), mice from all groups traveled an equal distance in the arena (*P* = 0.102), suggesting that viral constructs injected do not affect the overall locomotory behavior (Fig. S6 A). Next, three objects were placed in fixed locations (Fig. 5A, *middle*) and mice were allowed to explore them for 5min during each trial. Mice of all groups show a significantly decreased exploration of the objects across sessions (*F*_(2,154)_ = 18.31, *P* < 0.0001) (Fig. S6 B), signing the habituation of the exploratory response induced by exposure to novelty. In the last two trials, mice were tested in the novel object location task (Fig. 5A, *right*), in which one object (object A) was moved to a novel location while the other two objects (B and C) remained in their original position. These trials were used to evaluate the innate tendency of rodents to direct exploration toward objects that have been displaced. Lack of reaction to the spatial change has previously been used to assess cognitive impairments in mouse models of Alzheimer’s disease (Creighton et al., 2019). The reaction to the spatial change was expressed as percent contacts towards the displaced object and compared to the average of percent contacts with the two non-displaced objects. Both WT groups showed a significant reaction to spatial change (WT: n = 12, mean ± CI, 37.3 to 74.9 *P* = 0.022; WT UCP4: n = 11, mean ± CI, 36.31 to 67.96 *P =* 0.024) (Fig. 5, B and D) while mice in the 3xTg group did not exhibit exploration levels significantly (3xTg: n=17, mean ± CI, 30.57 to 54.64 *P =* 0.1207) above the chance level (33.3%). In contrast, overexpression of UCP4 in 3xTg mice, promoted a significant increase (3xTg UCP4: n=16, mean ± CI, 44.084 to 76.579, *P =* 0.003) in the exploration directed toward the displaced object (Fig. 5, C and D). These results indicate that uncoupling mitochondria in hippocampal astrocytes via exogenous expression of UCP4 improves reaction to spatial change that is impaired in aging 3xTg mice, in line with the observations shown above of rescued atrophy, dendritic architecture, neuronal excitability, and tissue metabolite profile.

**Fig. 5.**
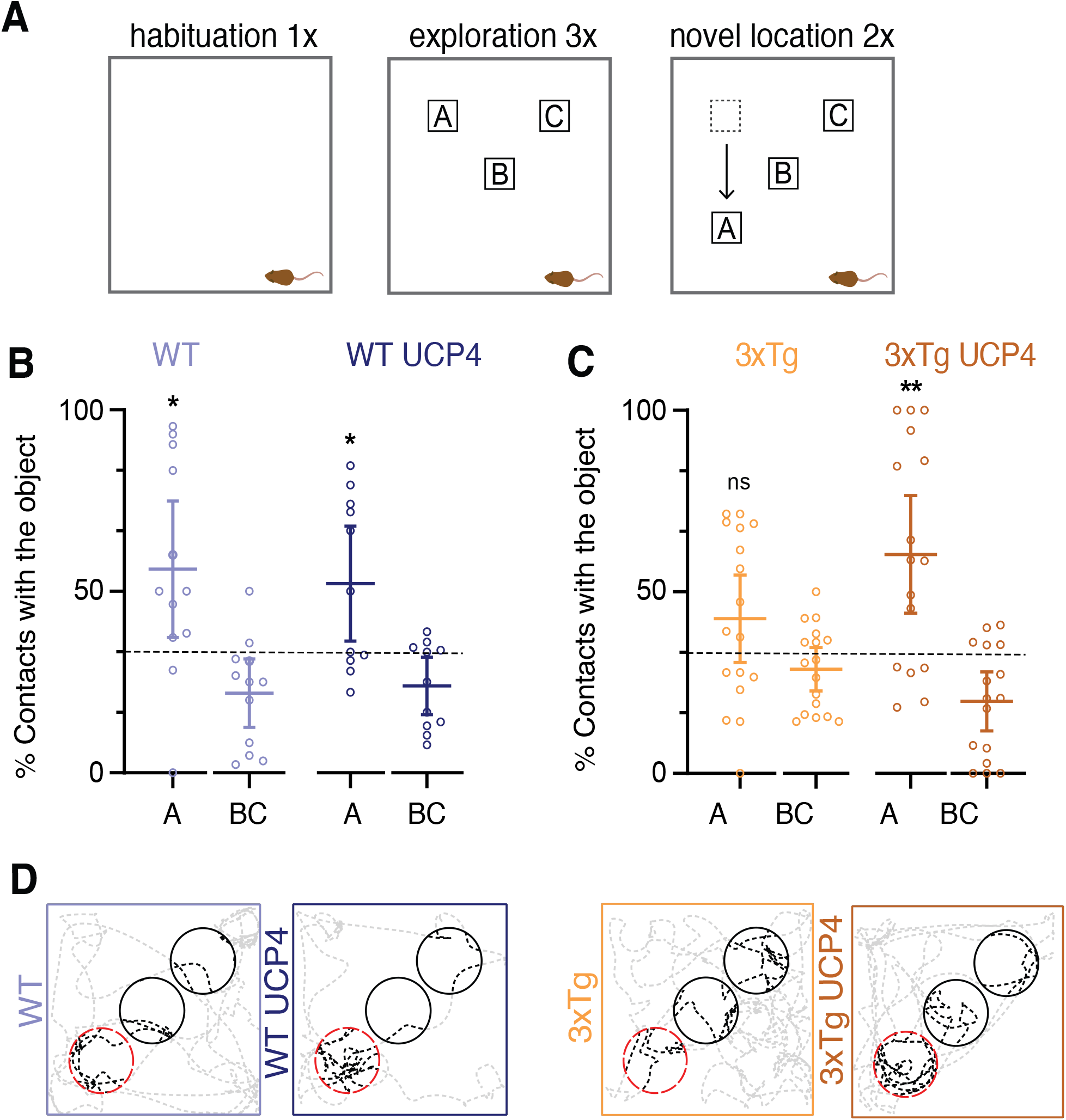
Mild mitochondrial uncoupling of astrocytes prevents spatial memory impairments of 3xTg-AD mice. (**A**) Schematic representation of the spatial recognition task. (WT: n=12, WT UCP4: 11, 3xTg: n=18, 3xTg UCP4: n=16). (**B**) and (**C**) Plots showing the reaction to spatial change expressed as the % of contact with the objects. A is the displaced object and BC is the average of the two objects that were not moved. Each symbol represents a different mouse. The dashed line represents the chance level (33.3%). (**D**) Representative tracking plots during the NOL trials. Black circles indicate the OEZ containing the objects that have not been moved (B and C). Red dashed circle shows the novel location zone of object A. Black dashed line represents the path followed by the mouse. In (**B**) and (**C**) one sample t test was performed. Error bars are mean ± CI. \**P* < 0.05; \*\*\*\**P* < 0.0001.

## Discussion

Current AD treatments attempt to counterbalance neurotransmitter disturbances or target aggregates of misfolded proteins, however so far with limited success. In this study, we selected a radically different approach which is of targeting astrocytes to better support neuronal function. We systematically investigated how *in vivo* uncoupling of astrocytic mitochondria impact phenotypes dysregulations ranging from metabolism to behavior occurring in a mouse model of AD. We showed that AD-associated altered metabolism, hippocampal atrophy, dendritic field shrinkage, burst frequency and AHP alteration of subicular neurons, and spatial memory decline are prevented by UCP4 overexpression in astrocytes.

We found that at an early stage of AD before the onset of neuropathology and memory decline, 3xTg mice present already a different metabolic profile compared to WT and 3xTg mice treated with UCP4. We show that lactate, other metabolites from glycolysis, TCA cycle, pyruvate metabolism, and histidine metabolism are significantly elevated in 3xTg mice compared to the other groups. These results could reflect an adaptative change to sustain neuronal hyperactivity observed in subicular neurons. Our observations are in line with a recent study (van der Velpen et al., 2019) showing higher concentrations in both plasma and CSF of TCA cycle intermediates in AD patients compared to control subjects. In addition, it has been shown that metabolites of glycolysis such as phosphoenolpyruvate, are increased in 3xTg male mice hippocampus compared to non-transgenic mice at 3 months of age (Dong & Brewer, 2019). However, we do not have the means to distinguish whether an increased level of certain types of metabolites results from upregulation of a specific metabolic pathway, or if it is due to an impairment of catabolic enzymes involved in the pathway. The increase in lactate observed in 3xTg mice could thus result from glycolysis upregulation or from impaired neuronal consumption where metabolites would not enter the TCA cycle and accumulate in the cell. Finally, while not addressing the mechanistic that keeps metabolites at basal level in AD mice overexpressing UCP4, we could speculate that astrocytic uncoupling enables 3xTg neurons to maintain physiological functioning, avoiding the buildup of oxidative stress linked with altered metabolism and may in addition, prevent the appearance of unbalanced expression levels of key enzymes of the metabolism.

At the structural level, MRI data show that UCP4 overexpression prevented the development of hippocampal atrophy in 3xTg-AD mice. The atrophy observed in 3xTg hippocampi, could partly be the consequence of dendritic branch loss and reduced arborization observed in 3xTg subicular neurons, also seen in different AD mouse models (Moolman et al., 2004; Šišková et al., 2014). These changes in dendritic morphology could arise from alterations in mitochondrial distribution, which was shown to be regulated in an ATP-dependent manner (Mironov, 2007). Thus, in a simplified scenario, the altered distribution of mitochondria across the neuron leads to their depletion from dendrites (Wang et al., 2009), diminished calcium clearance in spines, up-regulation of calcium signaling, increase in LTD and loss of spines resulting in atrophy of the dendritic tree (Lin & Koleske, 2010).

Truncated dendritic arborization observed in 3xTg subicular neurons, which is preserved by UCP4 treatment, can result in enhanced cellular excitability (Šišková et al., 2014) translated into an increase in AHP amplitude and burst frequency as seen in our results. These experimental observations are supported by our modeling data, in which the WT model with reduced morphology caused the transition from a single spike to bursting. Moreover, the WT model with increased SK and LVA conductance, simulating a 3xTg neuron, increased burst firing frequency. In shorter dendrites, the signal induced by synaptic stimulations can propagate more efficiently towards the axosomatic compartment without being diluted by backpropagation in more distal dendrites (Stuart et al., 1997). In parallel, a larger AHP could arise from exaggerated calcium release from the endoplasmic reticulum, for instance resulting from the upregulation of ryanodine receptors as described in 3xTg-AD mice (Stutzmann et al., 2006). It has been shown that SK2 channel activity is increased in CA1 neurons of 3xTg-AD mice (Chakroborty et al., 2012), along with the coupling between SK channels and ryanodine receptors (Stutzmann et al., 2006). We found evidence by enhancing SK channel activity, that the increase in AHP, in addition to be likely exacerbated by intracellular calcium, also results from an enhancement of SK channel activation. As SK protein levels have been reported to be unchanged in AD mice (Chakroborty et al., 2012), its increased activation could arise from a change in biophysical conformation increasing the conductance of the channel. These changes in excitability counterbalanced by astrocytic UCP4 overexpression, could contribute to synaptic plasticity alterations and subsequent AD-related network dysfunction, resulting in cognitive impairments.

Progressive memory loss is the hallmark of AD and is among the first symptoms reported by patients and their families. At first, only short-term memory is impaired followed by fragmentation of long term memory. In the early stages of AD, patients often suffer from spatial memory impairment, such as spatial disorientation, and tend to forget recently acquired information, *e.g.* fail to remember the location of an object soon after seeing it. This behavior involves spatial working memory that can be addressed in rodents with an object spatial recognition task (Dere et al., 2005). We show that astrocytic overexpression of UCP4 allows 3xTg mice to maintain a normal exploring behavior towards the object found in the novel location. The pattern of memory impairments in patients correlates with parameters defining both structural and functional brain integrity. Thus, as spatial memory is hippocampus-dependent (Broadbent et al., 2004), this statement fits our data that show correlation between hippocampal atrophy and poor spatial memory performance of AD mice, and conversely, normal memory performance when structural alteration of the hippocampus is prevented in mice overexpressing astrocytic UCP4.

In conclusion, we propose an innovative strategy targeting astrocytic mitochondria to preserve cellular and cognitive functions by tackling energy hypometabolism, a phenomenon occurring before neuropathology and memory decline in AD (Nunomura et al., 2001; Yao et al., 2009) and thus, providing strong potential for the development of efficacious therapies for AD and potentially for other brain pathologies.

## Supporting information

supplementary figures with legends

## Acknowledgments

Supported by the Synapsis Foundation Alzheimer Research Switzerland (JYC), and by the Blue Brain Project, a research center of the École polytechnique fédérale de Lausanne, EPFL, from the Swiss government’s ETH Board of the Swiss Federal Institutes of Technology (HM, MR). We thank A. Benechet and R. Colotti (IVIF, Unil-CHUV) for their help with MRI experiments. A. Arnaudon (BBP) for assistance with morphometrics packages. W. Van Geit (BBP) for the supervisory support. We thank J. Ivanisevic and colleagues of the Metabolomics Platform of the Faculty of Biology and Medicine at University of Lausanne for their expert handling of the metabolomic profiling. The behavioral experiments were run in the neuro-behavioral core of the Dept. of Fundamental Neurosciences, Univ. Lausanne. We thank M. Mameli (Univ. Lausanne) for his valued comments on the manuscript.

## Author Contributions

Conceptualization: NR, JYC, ABR; Methodology: NR, YB, LR, JYC; Investigation: NR, MR, MB; Visualization: NR, MR, LR; Funding acquisition: ABR, HM, JYC; Project administration: JYC; Supervision: JYC, NR; Writing – original draft: NR, JYC; Writing – review & editing: JYC, NR, MR, LR, ABR

## Data availability

The data underlying the study will be made available in the public repository Zenodo.org.

