## supplementary figures with legends for "Overexpression of UCP4 in astrocytic mitochondria prevents multilevel dysfunctions in a mouse model of Alzheimer’s disease"

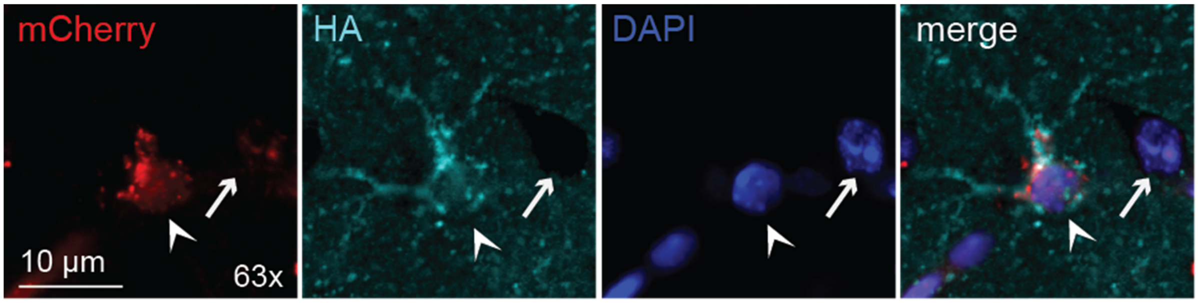


**Fig. S1. UCP4 expression at 4 months post-injection.** Confocal image of an astrocyte and a neuron in the CA1 region (highlighted with a white arrowhead and a white small arrow, respectively) showing HA-tag (cyan) fluorescent immunostaining and mCherry (red) plus DAPI (blue) revealing specific HA-tag expression in astrocyte only at 4 months in WT mice.

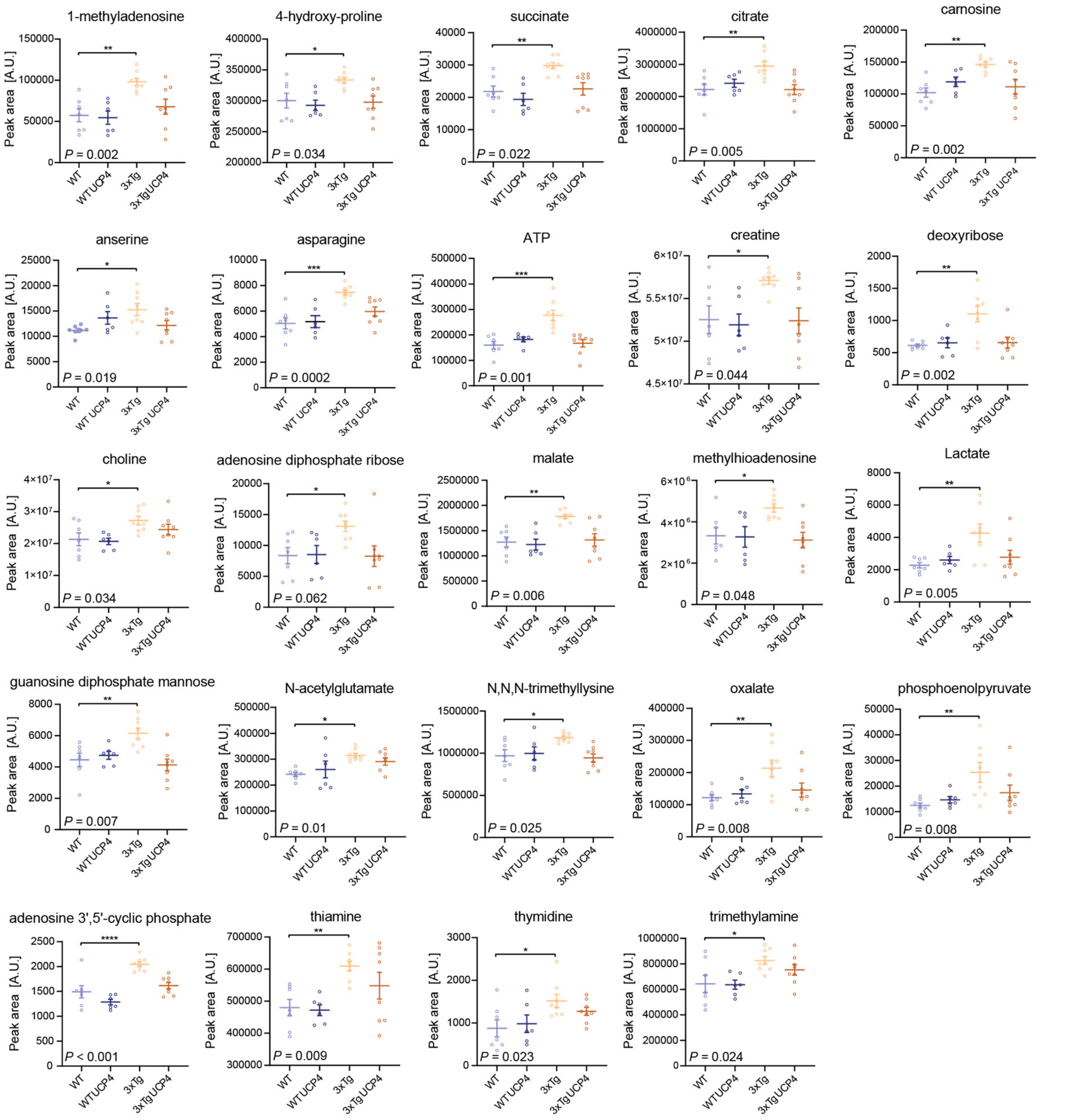


**Fig. S2. Metabolites that are altered in 3xTg mice but which are maintained to WT levels by overexpression of UCP4.** Plots of selected metabolites comparing hippocampal levels in WT (n=7), WT UCP4 (n=6), 3xTg-AD (n=8), and 3xTg-AD UCP4 (n=8) mice. Each dot represents a different specimen. One-way ANOVA followed by Tukey's post hoc test was performed, except for succinate, ATP, adenosine diphosphate ribose, methylhioadenosine and phosphoenolpyruvate a Kruskal-Wallis followed by a Dunn's post hoc test was done. *P <0.05; **P <0.01; ***P <0.001; ****P <0.0001.

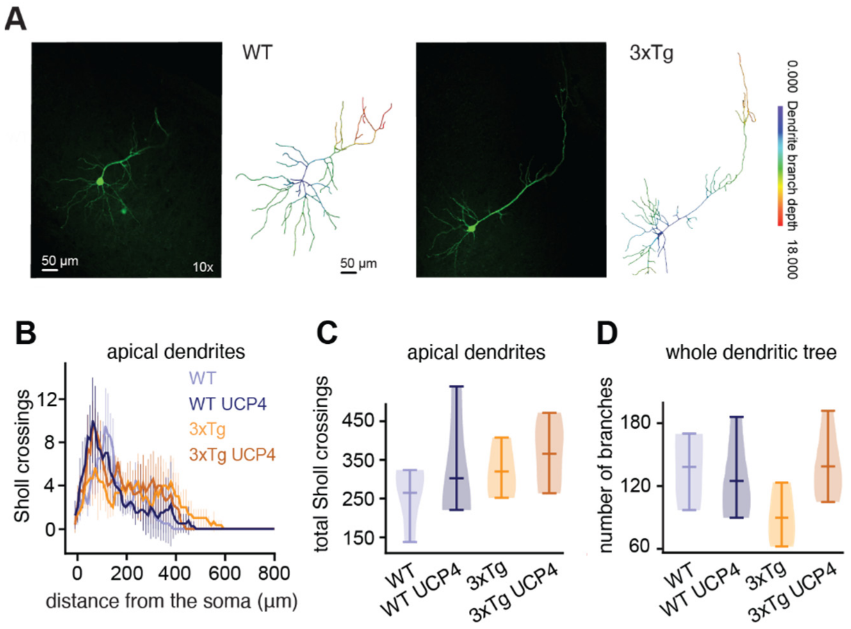


**Fig. S3. Sholl profiles and morphometrics of apical dendrites.** (**A**) Representative confocal images of WT (left) and 3xTg (right) subicular neurons filled with biocytin. The respective 3D reconstructions are shown with color-coded dendrite branch depth. (**B**) Sholl profile for each group, calculated using 5 μm increments (WT: n = 4, WT UCP4: n = 4, 3xTg: n = 3, 3xTg UCP4: n = 5). (**C**) Total number of Sholl crossing of apical dendrites for each group. No significant statistical difference was found between the groups. (**D**) Total number of dendritic branches for each group, calculated for each morphology. In **C** and **D** a non-parametric bootstrapping test was performed: no significant statistical difference was found between the groups. Bars on violin plots are mean ± CI.

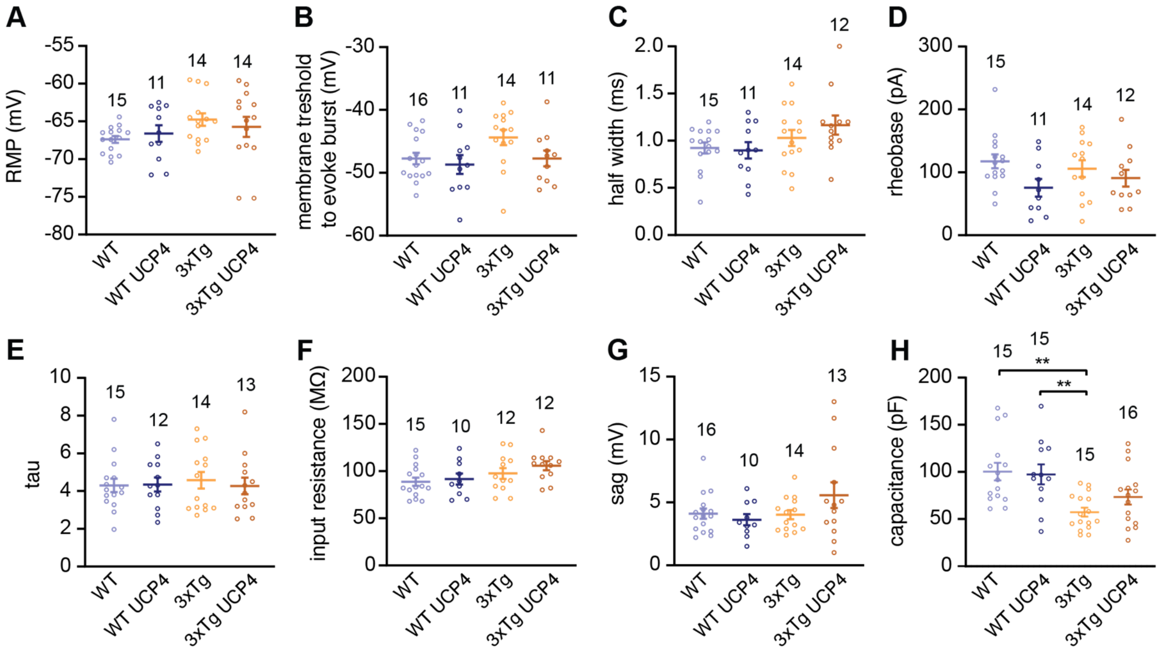


**Fig.S4. Intrinsic electrophysiological properties of subicular neurons are not altered by viral constructs.** Patch-clamp recordings of subicular neurons show no differences in intracellular properties, namely (**A**) resting membrane potential (RMP), (**B**) threshold to evoke burst, (**C**) AP half-width, (**D**) rheobase, (**E**) decay (tau), (**F**) input resistance, and (**G**) sag among groups, except for (**H**) capacitance. The numbers above each data set represent the number of neurons patched. In **A**-**D**, **F,** and **G** a one-way ANOVA followed by Tukey's *post hoc* test was performed, and in **E** and **H** a Kruskal-Wallis followed by a Dunn's post hoc test was done. Error bars are mean ± SEM. $^{**}P < 0.01$.

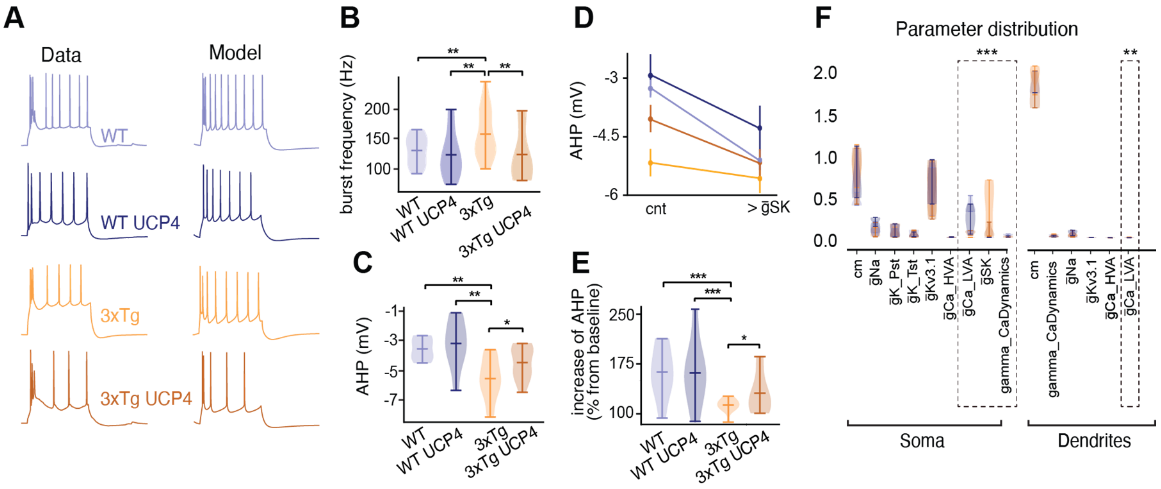


**Fig. S5. Generation and validation of subicular neuron models.** (**A**) Example traces of current clamp recordings (*Data*) and corresponding traces from neuronal models (*Model*). (**B**) Violin plot of burst frequencies in responses to simulated 200, 300 and 450pA current injections. Several models were considered for each group (WT: n=5, WT UCP4: n=5, 3xTg: n=7, 3xTg UPC4: n=5). (**C**) Violin plot of AHP in responses to pooled 200 and 450pA current injections. (**D**) AHP in control condition (cnt, same as in C), compared to a manually increased SK conductance (>ḡSK), across pooled models' responses to 200 and 450pA current injections. (**E**) Violin plot of AHP with increased ḡSK shown as percentage of control AHP amplitude. (**F**) Distribution of model parameters compared across groups of models (same color scheme as in (**A**)) Scale on the left: absolute value. Those include unitary capacitance (cm), sodium channel conductance (ḡNa), persistent potassium conductance (ḡK_Pst), transient potassium conductance (ḡK_Tst), potassium voltage gated channel conductance (ḡKv3.1) (Rudy & McBain, 2001), high voltage activated calcium channels conductance (ḡCa_HVA), low voltage activated calcium channel conductance (ḡCa_LVA) (Perez-Reyes, 2003), ḡSK, and gamma_CaDynamics, corresponding to the percentage of free intracellular calcium levels. Significant statistical difference between groups are highlighted by the dashed line rectangles. Scale on the left: absolute value. Bars on violin plots are mean ± CI. In (**B**), (**C**) and (**E**), nonparametric bootstrap test was performed. In (**D**) error bars represent mean ± SEM. In (**F**), Kruskal-Wallis, followed by a *post hoc* Bonferroni correction test was performed (somatic ḡCa_LVA (maximum conductance of low threshold calcium channels; KS: P = 3.9e-4, post hoc Bonferroni correction: 3xTg vs WT: P = 1.4e-7, 3xTg vs WT UCP4: P = 2.2e-5, 3xTg vs 3xTg UCP4: P = 1.2e-3, WT vs 3xTG-UCP4: P = 2e-3, alpha = 0.01), somatic ḡSK (maximum conductance of SK channel, KS: P = 4.5e-4, post hoc Bonferroni correction 3xTg vs WT: P = 2e-6, 3xTg vs WT UCP4: P = 5e-6, WT vs 3xTg UCP4: P = 4e-4, alpha = 0.01), somatic gamma_CaDynamics (percentage of free intracellular calcium; KS P = 5e-4, post hoc Bonferroni correction 3xTg vs WT: P = 1e-4, WT vs 3xTg UCP4 P = 8e-6, WT UCP4 vs 3xTg UCP4 P = 1.3e-4, alpha = 0.01) and apical ḡCa_LVA (KS: P = 2.9e-4, post hoc Bonferroni correction 3xTg vs WT: P = 5e-4, 3xTg vs WT UCP4 P = 3e 2, WT UCP4 vs 3xTg alpha = 0.01). **P* < 0.05; ***P* < 0.01; ****P* < 0.001. Bars on violin plots are mean ± CI.

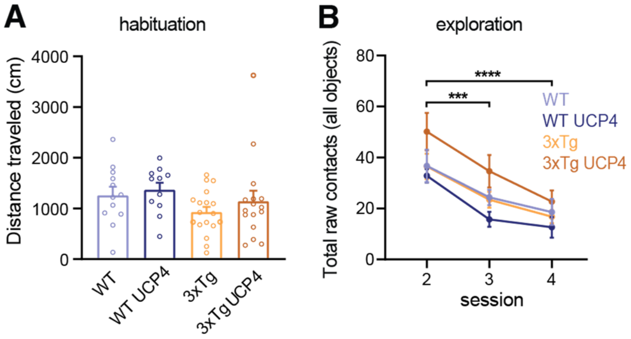


**Fig. S6. Viral vectors do not affect overall locomotory behavior**. (**A**) Distance traveled by the mice during the habituation trial. (WT: n=12, WT UCP4: 11, 3xTg: n=18, 3xTg UCP4: n=16). (**B**) Overall number of contacts with the 3 objects across sessions. Data are the average contact number with objects (same n per group as in **A**). In (**A**) a Kruskal-Wallis followed by a Dunn's test was performed. In (**B**) a two-way ANOVA followed by Tukey's *post hoc* test was performed, and the error bars are mean ± SEM. ***$P < 0.001$; ****$P < 0.0001$.
